# TBX2 driven switch from Androgen Receptor to Glucocorticoid Receptor signaling confers therapeutic resistance in Prostate Cancer

**DOI:** 10.1101/2023.05.07.539754

**Authors:** Sayanika Dutta, Girijesh Kumar Patel, Hamed Khedmatgozar, Daniel Latour, Manisha Tripathi, Srinivas Nandana

## Abstract

Recent studies have highlighted that androgen receptor (AR) signaling can be bypassed via activation of the glucocorticoid receptor (GR), and that this bypass drives enzalutamide resistance in advanced prostate cancer (PCa). However, the molecular mechanism(s) that drive the switch from AR to GR signaling remain unknown. We have previously reported that TBX2, a developmental T-box transcription factor (TF), is over-expressed in castrate resistant prostate cancer (CRPC) and that TBX2 drives the CRPC phenotype via cell-intrinsic and exosome-mediated paracrine modes. Our current study demonstrates that TBX2, a TF with known repressor and activator functions, may be the molecular switch that represses AR on one hand while activating GR expression on the other to drive CRPC. Mechanistically, our studies revealed a two-tiered mechanism of AR repression by TBX2 wherein TBX2 directly binds to the promoters of AR and GATA2, an AR coregulator, thereby resulting in the repression of AR as well as GATA2. Conversely, our results demonstrate that TBX2 mediates increased expression of GR via directly binding to the GR promoter, and through TBX2-GR functional protein-protein interaction. Our results demonstrate that the TBX2 driven switch from AR to GR signaling results in enzalutamide resistance since GR inhibition in the context of TBX2 over-expression attenuates enzalutamide resistance. Further, we present evidence that SP2509 based allosteric inhibition of Lysine Specific Demethylase 1 (LSD1), a protein that interacts with TBX2 as part of the Co-repressor of RE1-Silencing Transcription Factor (COREST) complex, is able to disrupt TBX2-GR interaction. Taken together, our study has identified TBX2 as the molecular switch that drives AR to GR signaling and thereby confers enzalutamide resistance in CRPC. Furthermore, our study provides key insights into a potential therapeutic strategy of targeting the AR to GR switch wherein SP2509-based allosteric inhibition of TBX2-LSD1 could be harnessed to target the TBX2-GR interaction, thereby resulting in the inhibition of enzalutamide resistance in CRPC.

## Introduction

Androgen deprivation therapy (ADT) that includes androgen receptor signaling inhibitors (ARSIs) forms the cornerstone of treatment of advanced prostate cancer (PCa) ^1–3^. Following a period of response, patients inevitably develop resistance to ADT when the disease progresses to castration resistant prostate cancer (CRPC) ^1–4^. CRPC is associated with heterogenous expression of AR (AR positive and AR low/negative CRPC) ^5–7^. In recent years, the use of more potent ADTs like enzalutamide has led to significant improvements in patient survival, however, the increased selection pressure on the PCa cells leads to clinically unintended consequences wherein PCa is driven towards AR signaling bypass mechanisms ^7–11^. More recently, it has been reported that resistance to enzalutamide can be driven by upregulation of the glucocorticoid receptor (GR) ^9, 11–14^, a steroid nuclear receptor whose DNA binding domain shares a high level of similarity with the DNA binding domain of AR ^8–11, 13^. As a result, both AR and GR share significant overlaps in their cistromes and transcriptomes ^8–10, 13, 15^, and these reports have led to a hypothesis in the field that the signaling bypass from AR to GR drives CRPC ^7–11, 16^. However, the molecular mechanisms that drive the AR-GR signaling bypass remain to be identified.

Recent reports have demonstrated that TBX2 physically interacts with the Co-repressor of RE1-Silencing Transcription Factor (CoREST) complex that includes Lysine Specific Demethylase 1 (LSD1) and that SP2509, an allosteric LSD1 inhibitor, was able to disrupt TBX2 interaction with the CoREST complex including LSD1 resulting in and diminished cancer cell growth and survival ^17^.

Previous reports ^18, 19^ from our lab have demonstrated that TBX2, a T-box family of transcription factor (TF) ^20^: a) drives the PCa bone metastatic cascade^18, b^) is over-expressed in models of human CRPC progression^18^, and c) mediates propagation of the CRPC phenotype via cell-intrinsic and exosome-mediated paracrine modes^19^. In accordance with our findings, a recent report identified TBX2 and GR as two of the four TFs that drive enzalutamide resistance ^21^. In the current paradigm-shifting study, we have identified TBX2 as the molecular switch that drives AR to GR signaling bypass thereby conferring enzalutamide resistance. Further, our studies provide key insights into a potential therapeutic modality of targeting the bypass from AR to GR signaling through disruption of the TBX2-LSD1 and TBX2-GR protein-protein interactions. Given that systemic ablation of GR has deleterious effects on PCa treatment ^12, 22^, our study may offer an indirect and potent molecular mechanism for GR inhibition that could restore enzalutamide sensitivity in CRPC.

## Materials and Methods

### Cell lines and growth conditions

The human PCa cell lines-PC3, C4-2B, LNCaP ^23, 24^, 22Rv1 were procured and maintained at 37°C in humidified 5% CO2 incubator using either Dulbecco’s Modified Eagle Media (DMEM, corning Cat # 10013CV) or Roswell Park Memorial Institute media −1640 (RPMI-1640 corning Cat #10040CV) supplemented with 10% FBs and 1% penstrep. The PCa cells were received from Dr. Leland W. K Chung, Uro-Oncology research program, Department of Medicine, Cedars-Sinai Medical Center, Los Angeles, California, USA. The cell lines used in this study were evaluated for mycoplasma contamination in house.

### RNA isolation, cDNA synthesis and quantitative Real Time RT-PCR (qRT-PCR)

Total RNA was isolated from PCa cells using the RNAse easy mini kit (Qiagen Inc, Valencia, CA, USA) by following the manufacture’s protocol. This was followed by cDNA synthesis and qRT-PCR as described before ^19^. Quantstudio 12k Flex qRT-PCR platform was used to perform the qRT-PCR. The relative amount of mRNA expression was normalized with β-actin or GAPDH.

### Cell lysate preparation and Western blot analysis

Total protein lysate from PCa cells were isolated followed by protein quantification as described previously ^19^. Equal amounts of protein were resolved onto SDS-PAGE gel and electroblotted onto PVDF membrane (IPVH00005)). The membranes were blocked with 5% BSA in TBST and incubated with respective primary antibodies overnight at 4°C. After washing, the membranes were probed with HRP-conjugated secondary antibodies (rabbit/mouse, CST). The bands were visualized using west pico chemiluminescent kit (#34580, Thermo Fisher Scientific, Rockford, IL) under chemi-doc touch imaging system. β-actin was used as a loading control.

### Chromatin immunoprecipitation (ChIP) assay

Using *in silico* analysis tools like JASPAR ^25^ and published literature we predicted the T-box consensus binding sequences on AR, GATA2 and GR promoter. Chromatin immunoprecipitation (ChIP) assays were performed as described in previous literature ^19^. The DNA was purified using PCR purification kit (Qiagen, Hilden, Germany) and amplified by qRT-PCR.

### Site Directed Mutagenesis (SDM)

AR promoter upto 4000bp upstream of transcription start site (TSS) was cloned in pMCS-Cypridina Luc Vector (#16149) and was procured from Custom DNA Constructs, Islandia, NY. Site directed mutagenesis (SDM) was performed by mutating using base substitution on the two TBX2 binding sites (−82bp, −3598bp) ^26^. The mutagenic primers were designed using QuickChange primer design program at www.agilent.com/genomics/qcpd according to the guidelines mentioned in the manufacturer’s protocol (QuickChange Lightning Site-Directed Mutagenesis kit, Cat# 210518, Agilent Technologies). This was followed by mutant strand synthesis and transformation was performed as mentioned in the manufacturer’s protocol. The transformed plates were incubated at 37° for >16 hours and mutant colonies were observed the next day. The mutant colonies were further grown in bulk in LB-amp broth and plasmid isolation was performed following manufacturer’s protocol.

### Luciferase reporter assay

Luciferase reporter assay was performed using Pierce^TM^ Cypridina-Firefly Luciferase Dual Assay Kit (Cat#16183). After transfection of the mutated cypridina luc reporter plasmid and red firefly luc control plasmid (cat#16156), the cells were incubated for 16-72 hours at 37°C in 5%CO2 in a cell culture incubator. Luciferase reporter assay was performed according to manufacturer’s protocol. The reading of cypridina luciferase signal was normalized to control red firefly luciferase signal and the fold change in luciferase activity was calculated.

### Plasmid isolation, transduction and modulation of TBX2 and GR using sh-RNA/ si-RNA

The custom constructs for sh-TBX2 and its non-targeting scrambled RNA duplex siRNA control (NTSCR) was procured from sigma Aldrich (RNAi single clones, Millipore sigma, Burlington, MA). These lentiviral constructs along with packaging plasmid (pCMV delta R8.2 dvpr, addgene#8455) and envelope plasmid (pCMV-VSV-G, Addgene#8454) were used for lipofectamine 2000 mediated transfection of 293FT cells to produce lentiviral pseudo typed particles. The supernatant containing viral particles were collected post 36h. Infection with viral particles was repeated three times and selection was performed with neomycin at 500 µg/ml concentration. The si-RNA mediated knockdown of GR performed using si-GR construct (SCBT # SC35505) and control si-RNA (SC37007) were purchased from Santacruz biotechnology. The si-RNA transfection was performed in LNCaP^TBX2OE^ cells following manufacturers protocol. Following transfection, the cells were incubated for 18-24h after which fresh 1X normal growth serum was added along with enzalutamide treatment and the cells were assayed for cell viability after 72 h of treatment.

### Cell viability assay

Cell viability assay was performed according to standard manufacturer’s protocol. Briefly about 5*10^3^ cells/well were seeded in 96-well plates in five replicates. This was followed by different treatment conditions. Cell proliferation was examined at 48, 96, and 72 h using WST-1 reagent from Roche Applied Science (Indianapolis, IN). Absorbance was measured at 450nm before saturation on iMARK plate reader (Bio-Rad). For background control, 1XWST-1 was added in empty wells in triplicate and absorbance was subtracted and relative percentage of cell proliferation was calculated.

### Co-immunoprecipitation (Co-IP)

Co-IP was performed as described previously ^27^ with minor modifications. Immunoprecipitation was set using anti-TBX2 antibody (SC-514291, SCBT, Dallas, TX) for 2hours at 4°C and this was followed by precipitation of the protein-antibody complex with dynabeads protein G (Invitrogen, #300D, Carlsbad, CA, USA) for 2h in 4°C on rotator. The immunoprecipitated samples were washed with lysis buffer and eluted in 1X gel loading buffer.

### Statistical analysis

All the experiments were executed at least three times in a biological replicates and data expressed as mean ± SD. Wherever appropriate, data were subjected to unpaired two-tailed Student’s t-test for comparison of two groups and one-way ANOVA for three groups or more. Differences were considered statically significant when *p ≤ 0.05. Statistical calculations were executed using GraphPad prism8 where *≤ 0.05, ** p≤0.01, *** p≤0.001, **** p≤0.0001.

## Results

### Blocking TBX2 in human PCa results in increased AR and AR target genes

Our previous study using xenografts of isogenic human PCa cell lines that mimic human PCa progression from adenocarcinoma (LNCaP) to CRPC (C4-2) to bone metastatic CRPC (C4-2B) demonstrated an increase in TBX2 expression with the disease progression^18^. Therefore, we sought to investigate a potential relationship between TBX2 and AR ^18^. An analysis of 488 human PCa samples in cBioportal ^28, 29^ revealed a negative correlation between TBX2 and AR (Spearman −0.31, p=1.64e-12) (**Fig. 1A**) indicating an inverse association between TBX2 and AR. Therefore, we sought to investigate the relationship between TBX2 and AR signaling using genetic modulation approaches for TBX2. Since previous reports including our studies have shown that TBX2 predominantly acts as a repressor of gene expression, we utilized the dominant negative (DN) approach of blocking TBX2 wherein the repressor domain of the protein has been deleted ^30^. Using a converse approach, we over-expressed (OE) TBX2 in LNCaP cells that express low endogenous TBX2. Utilizing an unbiased approach of performing RNA-seq in TBX2 blocked PC3 human CRPC cells, a cell line that does not express the AR at either the mRNA or protein levels ^31^, we observed a dramatic (60-fold) increase in AR mRNA expression in PC3^TBX2DN^ cells when compared with the PC3^Neo^ control cells (**Fig. 1B**). In addition, multiple AR target genes were significantly upregulated (TMPRSS2, PMEPA1, STEAP4) ^32^ (**Fig. 1B**). Validation of the RNA-seq at mRNA and protein levels revealed increased AR expression when TBX2 is blocked (PC3^TBX2DN^ and C4-2B^TBX2DN^ compared with the respective controls i.e. PC3^Neo^ and C4-2B^Neo^ and conversely, decreased AR expression in LNCaP^TBX2OE^cells when compared with the LNCaP^Neo^ control (**Figs. 1C, D)**. In addition, some of the known AR downstream targets (TMPRSS2, NKX 3.1 and FKBP5 ^33^ ^34^ showed a negative association with TBX2 expression in LNCaP, C4-2B and PC3 cells (**Fig. 1E**). As an additional method of validation, we utilized the sh-RNA method to knock down TBX2 expression in PC3 cells and observed similar results that showed upregulation of AR in PC3^shTBX2^ compared with PC3^NTSCR^ control (**Fig. 1F**). Taken together, these results pointed to increased AR expression as well as increased AR target genes upon blocking TBX2 in human PCa.

**Fig. 1:**
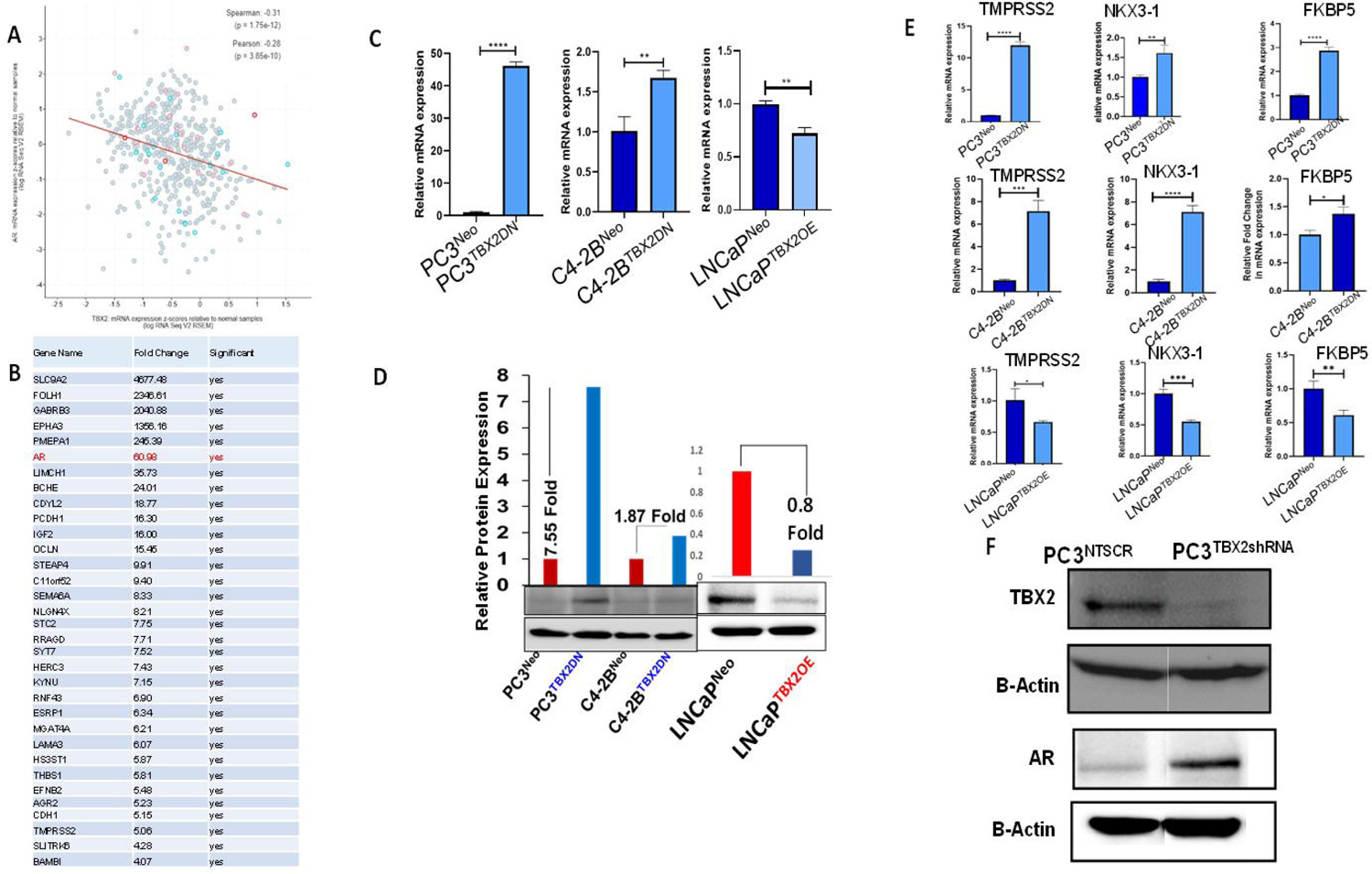
Blocking TBX2 in human PCa results in increased AR and AR target genes. **A)** Human PCa samples (n=488) from cBioportal showing a negative correlation between TBX2 and AR; **B)** RNA-seq analysis in PC3^TBX2DN^ compared with PC3^Neo^ control showing a significant increase in the expression of AR (60-fold) and AR target genes; **C)** q RT-PCR analysis showing increased AR mRNA expression in PC3^TBX2DN^ and C4-2B^TBX2DN^ when compared with the respective PC3^Neo^ and C4-2B^Neo^ controls, and conversely a decreased AR in LNCaP^TBX2OE^ cells when compared with LNCaP^Neo^ control; **D)** Western blot analysis showing increased AR protein expression in PC3^TBX2DN^ and C4-2B^TBX2DN^ cells when compared with the respective PC3^Neo^ and C4-2B^Neo^ controls, and conversely decreased AR protein in LNCaP^TBX2OE^ cells when compared with LNCaP^Neo^ control; **E)** qRT-PCR analysis showing increased mRNA expression of some of the known AR target genes in PC3^TBX2DN^ and C4-2B^TBX2DN^ cells when compared with the respective PC3^Neo^ and C4-2B^Neo^ controls, and conversely decreased expression of the AR target genes in LNCaP^TBX2OE^ when compared with LNCaP^Neo^ control; **F)** Western blot analysis showing decreased TBX2 and increased AR protein expression in PC3^TBX2shRNA^ cells when compared with the PC3^NTSCR^ control. Data represent the average of triplicates ± S.D; Student’s unpaired 2-tailed *t*-tests were performed to compare the two groups **, *p*≤ 0.01; ***, *p*≤0.001; and ****,*p*≤0.0001.

### TBX2 directly represses AR transcription

Since our genetic modulation studies showed a negative association between TBX2 and the AR, including the AR target genes, we next investigated the possibility if the TBX2 TF repressed AR via direct transcriptional regulation. Based on the consensus T-Box binding sequences that have been previously reported ^35^, *in silico* analysis revealed multiple sites on the AR promoter region to which TBX2 could potentially bind (**Fig. 2A**). Chromatin immunoprecipitation (ChIP) followed by RT-PCR and quantitative real-time RT-PCR (qRT-PCR) analyses revealed a 5-fold and 10-fold enrichment of TBX2 binding at the −82 bp and −3598 bp upstream of the AR transcription start site (TSS) respectively, indicating TBX2 binding to these sites (**Figs. 2B, 2C**). To further confirm AR transcriptional repression by TBX2, we performed site directed mutagenesis (SDM) experiments using base substitution wherein we mutated the two TBX2 binding sites of the AR promoter (−82 bp and −3598 bp) that we had previously identified (**Fig. 2D**), followed by luciferase reporter assays. The mutated AR-luciferase plasmids were transfected in to wildtype PC3 cells and luciferase reporter assays were performed. These experiments revealed a significant increase in the AR-luciferase activity for each of the mutant sites (**Fig. 2E**). Taken together, these results that show that: a) blocking/knock-down of TBX2 in human PCa cells markedly increases the AR, and b) mutating TBX2 binding sites on the AR promoter increases AR transcription – strongly demonstrates that TBX2 directly represses AR transcription.

**Fig. 2:**
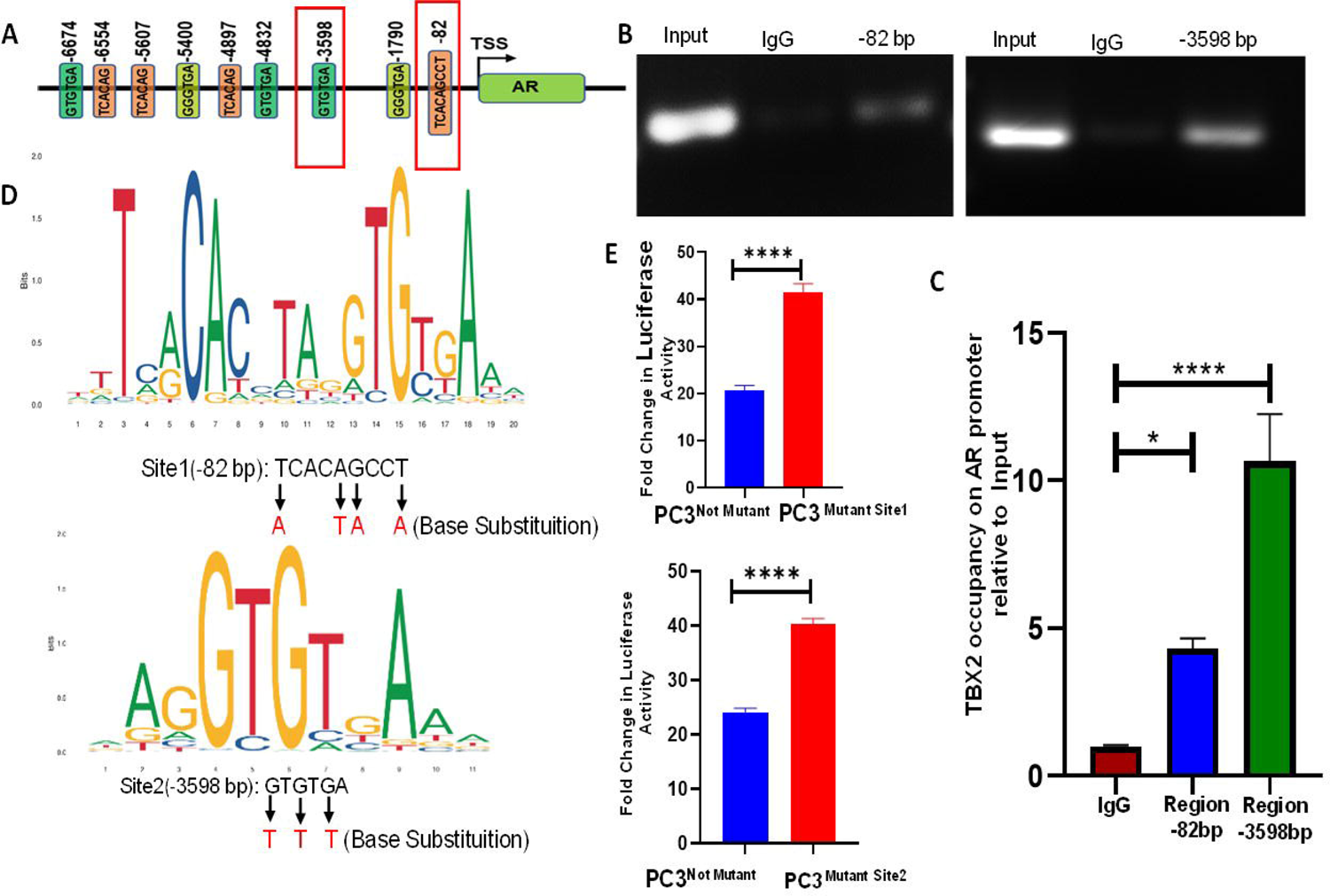
TBX2 binds to the AR promoter and represses AR transcription. **A)** *In silico* analysis of the AR promoter showing predicted TBX2 binding sites on the AR promoter; **B)** Chromatin immunoprecipitation (ChIP) demonstrating TBX2 binding at the −82 bp and −3598 bp upstream of the transcription start (TSS) site on the AR promoter, **C)** Quantitation of the ChIP assay using qRT-PCR analysis shown in B demonstrating a 5-fold and 10-fold enrichment of TBX2 binding normalized to the input on the −82 bp and −3598 bp binding sites on the AR promoter. One way ANOVA was performed (n=3), \*\*\**p*<0.001; ****, *p*<0.0001, **D)** JASPAR based depiction of the two consensus T-box binding sequences on the AR promoter. Arrows indicate mutated base pairs in the mutant site1 (−82 bp) and mutant site2 (−3598 bp) on the AR promoter; **E)** Quantitation of luciferase reporter assay in PC3 wildtype cells showing increased luciferase reporter gene activity in each of the two mutant sites i.e. mutant site1 (−82 bp) and mutant site2 (−3598 bp) normalized to the control. Data represent the average of triplicates ± S.D; Student’s unpaired 2-tailed *t*-tests were performed to compare the two groups **, *p*≤ 0.01; ***, *p*≤0.001; and ****, *p*≤0.0001.

### TBX2 repression of AR is additionally mediated through direct repression of GATA2, an AR coregulator

Previous studies have reported that GATA2 is a crucial coregulator of AR signaling that is necessary for the optimal transcriptional activity of AR ^36–39^. Since our RNA-seq data (**Fig. 3A)** and validation of RNA-seq at mRNA and protein levels (**Figs. 3 B, C, D)** pointed to increased GATA2 expression upon blocking TBX2 in PC3 human PCa cells, we therefore investigated if GATA2 is a mediator in TBX2 repression of AR. Further, *in silico* analysis identified multiple T-box binding sites on the GATA2 promoter to which TBX2 could potentially bind (**Fig. 3E**). ChIP assay followed by RT-PCR and qRT-PCR analyses revealed an 8-fold and 12-fold enrichment of TBX2 binding at the −510 bp and −460 bp upstream of the GATA2 TSS respectively, indicating TBX2 binding to these sites (**Figs. 3F, G**). Further, previous reports have identified that GATA2 binds at −4.6 kb on the AR promoter and activates its transcription ^39^. Therefore, we endeavored to determine if blocking TBX2 would drive an increase in GATA2 activation of the AR. ChIP analysis revealed that blocking TBX2 in PC3 human PCa cells (PC3^TBX2DN^) resulted in 10-fold enhanced GATA2 occupancy at binding site that is −4.6 kb upstream of the TSS of the AR (**Fig. 3H**). Taken together, our results strongly point to a two-tiered mode of AR repression by TBX2, i.e. by direct repression of: a) AR, and b) GATA2, an AR co-activator - thereby resulting in an enhanced repression of AR transcription.

**Fig. 3:**
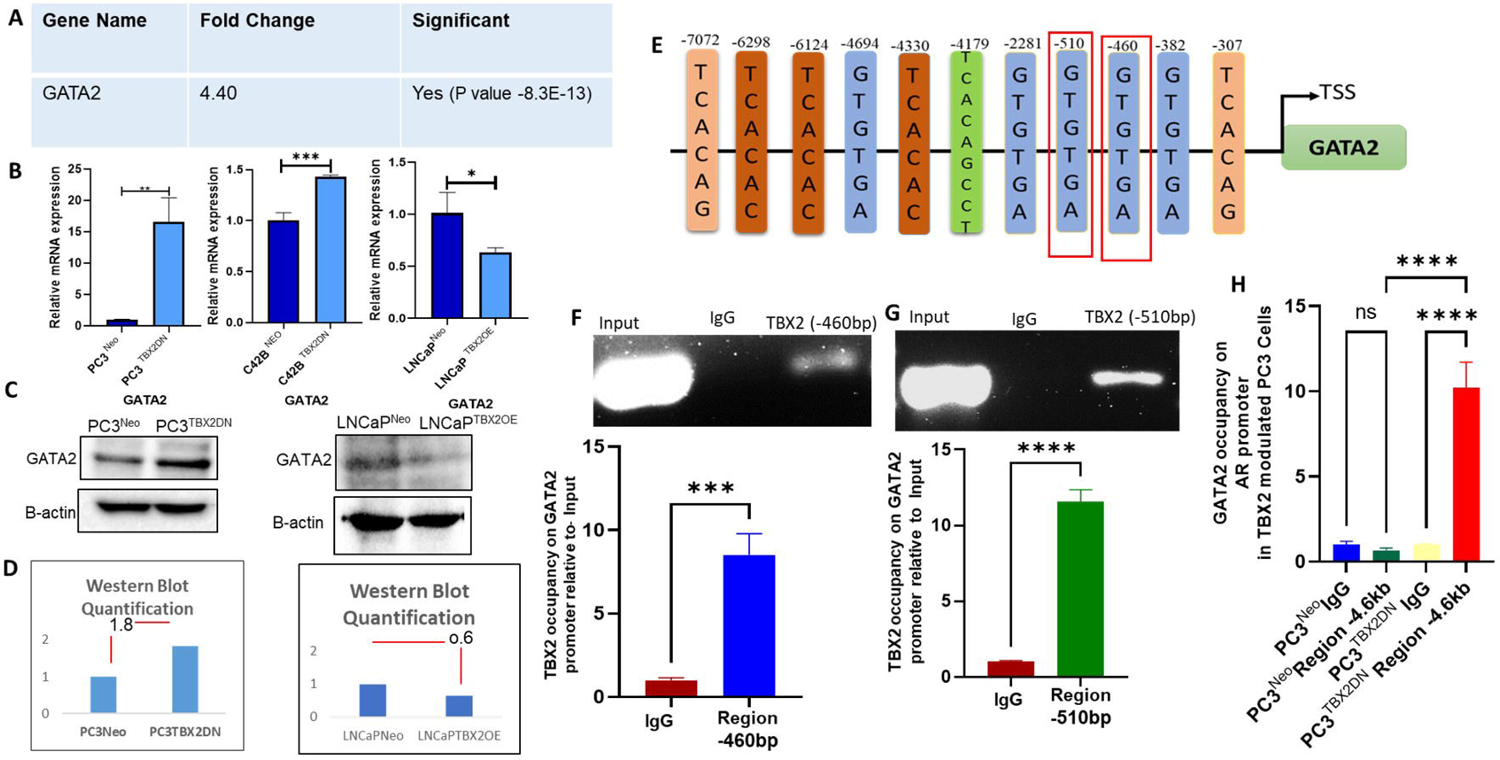
TBX2 represses AR by directly binding to the promoter of GATA2, an AR-coregulator. **A)** RNA-seq analysis in PC3^TBX2DN^ cells compared with PC3^Neo^ control showing a significant increase in the expression of GATA2 (4.4-fold, P value −8.3E-13); **B)** qRT-PCR analysis showing increased GATA2 mRNA expression in PC3^TBX2DN^ and C4-2B^TBX2DN^ when compared with the respective PC3^Neo^ and C4-2B^Neo^ controls, and conversely decreased GATA2 expression in LNCaP^TBX2OE^ cells when compared with LNCaP^Neo^ control; **C)** Western blot analysis showing increased GATA2 protein expression in PC3^TBX2DN^ cells when compared with the respective PC3^Neo^ control, and conversely decreased GATA2 protein in LNCaP^TBX2OE^ when compared with LNCaP^Neo^ control; **D)** Quantitation of the Western blot in C; **E)** *In silico* analysis of the GATA2 promoter showing the predicted TBX2 binding sites; **F, G)** ChIP assay followed by qRT-PCR showing 8-fold and 12-fold enrichments of TBX2 occupancy on the GATA2 promoter at the −460 bp and −510 bp respectively normalized to input in C4-2B cells. Data represent the average of triplicates ± S.D; Student’s unpaired 2-tailed *t*-tests were performed to compare the two groups **, *p*≤ 0.01; ***, *p*≤0.001; and ****,*p*≤0.0001; **H)** ChIP qRT-PCR in PC3^TBX2DN^ cells showing significant enrichment (10-fold) of GATA2 on the AR promoter at −4.6kb upstream of TSS normalized to input. One way ANOVA was performed where (n=3), \*\*\**p*<0.001;****,*p*<0.0001

### TBX2 is associated with increased GR expression in PCa

AR and glucocorticoid receptor (GR) belong to the steroid nuclear-receptor superfamily with highly conserved DNA binding domains ^8–10, 13^. Recent evidence has pointed to GR upregulation as a clinically unintended consequence of blocking AR in PCa, particularly with the use of 2^nd^ generation ADTs such as enzalutamide ^8–11, 13^. Studies have pointed to the significant overlaps in the cistromes and transcriptomes of AR and GR as the molecular basis for the re-establishment of an AR-like response in enzalutamide therapy ^8–11, 15^. Further, a recent report identified GR and TBX2 among the four TFs that drive enzalutamide resistance in human CRPC^21^. Therefore, we asked if TBX2 is associated with upregulation of GR in PCa. The cBioportal human database ^28, 29^ revealed a positive correlation between TBX2 and GR expression in 531 human PCa patient samples (Spearman 0.61, p=4.59e-18) (**Fig. 4A**). This observation was reinforced by a positive association between TBX2 and GR in human PCa cell lines genetically modulated for TBX2. Blocking TBX2 expression in C4-2B and 22Rv1 human CRPC cell lines significantly reduced GR expression; while the converse approach of TBX2OE in LNCaP^TBX2OE^ human PCa cells resulted in increased GR expression both at the mRNA and protein levels compared to control LNCaP^Neo^ (**Figs. 4B, C**). Taken together, our data shows that TBX2 is positively associated with GR expression in CRPC.

**Fig. 4:**
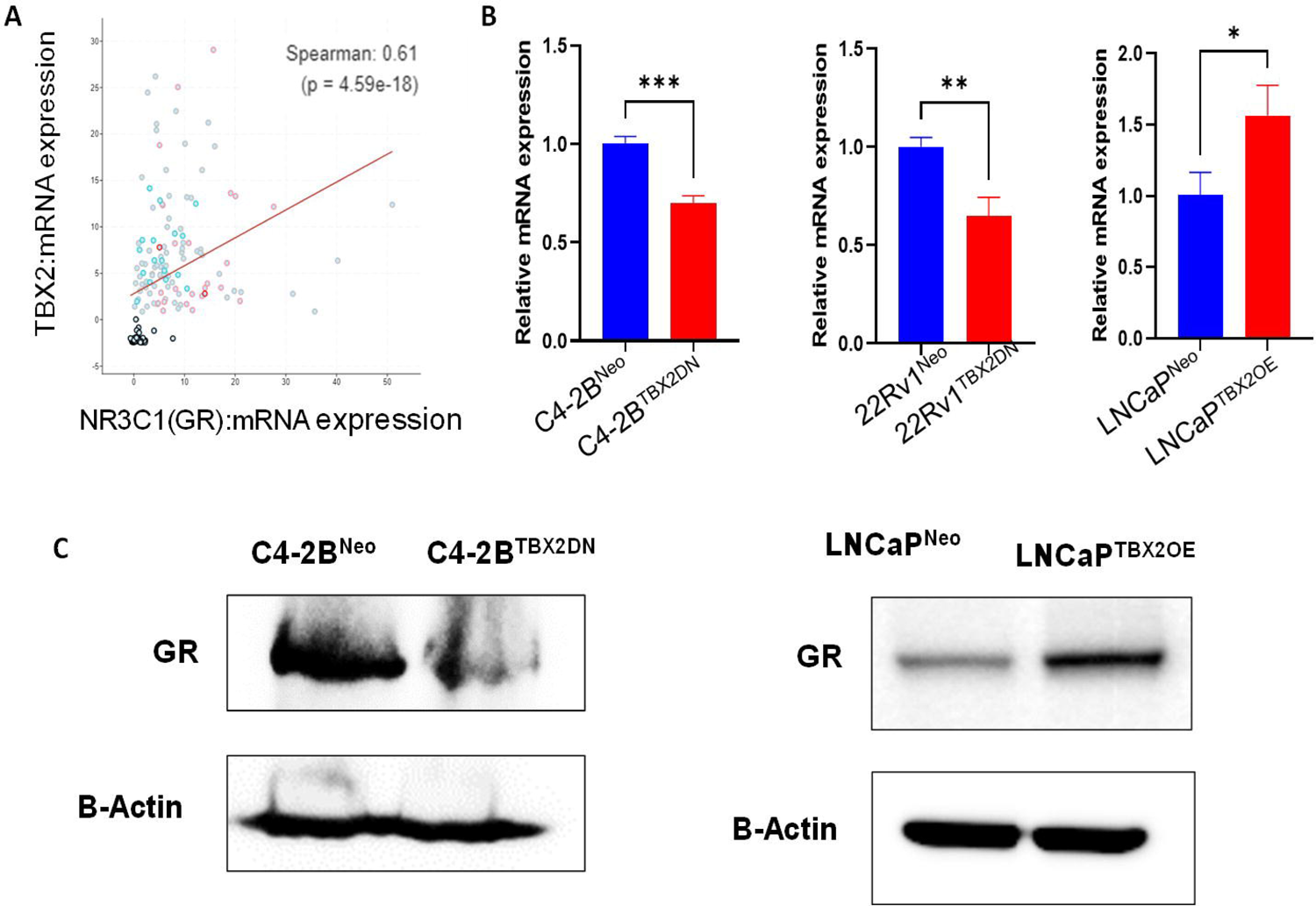
TBX2 is associated with increased GR expression in PCa. **A)** Human PCa samples (n=531) from cBioportal showing a positive correlation of TBX2 with GR (Spearman 0.61, p=4.59e-18); **B)** qRT-PCR analysis showing decreased GR mRNA expression in C4-2B^TBX2DN^ and 22Rv1^TBX2DN^ when compared with the respective C4-2B^Neo^ and 22Rv1^Neo^ controls, and conversely an increased GR expression in LNCaP^TBX2OE^ cells when compared with LNCaP^Neo^ control; Data represent the average of triplicates ± S.D; Student’s unpaired 2-tailed *t*-tests were performed to compare the two groups **, *p*≤ 0.01; ***, *p*≤0.001; and ****,*p*≤0.0001.; **C)** Western blot analysis showing decreased GR protein expression in C4-2B^TBX2DN^ cells when compared with the respective C4-2B^Neo^ control, and conversely increased GR protein in LNCaP^TBX2OE^ when compared with LNCaP^Neo^ control.

### TBX2 activates GR through transcriptional activation and protein-protein interaction

Thus far, our results point to a positive correlation between TBX2 and GR expression in CRPC. Next, we interrogated TBX2’s ability to directly activate the GR. Interestingly, T-box family members have been shown to either activate or repress gene expression ^40^. A previous report ^18^ from our lab demonstrated the ability of TBX2 as an activator of gene transcription. *In silico* analysis of the GR promoter revealed the presence of T-box consensus binding sites (**Fig. 5A**). We next performed ChIP followed by qRT-PCR analyses that revealed the presence of two TBX2 binding sites on the GR promoter, one at −710 bp and the other at −4854 bp upstream of the TSS, with significant enrichment of 12 and 8 folds respectively (**Fig. 5B**). These results pointed to GR transcriptional activation by TBX2. Further, in addition to direct transcriptional activation through promoter binding, TFs can also physically interact with proteins resulting in their activation ^41^. We next investigated if TBX2 interacts with the GR via a protein-protein interaction. A co-immunoprecipitation (CoIP) assay using the 22Rv1 and C4-2B human CRPC cell lines revealed a physical interaction between TBX2 and GR (**Fig. 5C**). These results demonstrate a dual mode of TBX2 regulation of the GR, i.e. through transcriptional activation and protein-protein interaction.

**Fig. 5:**
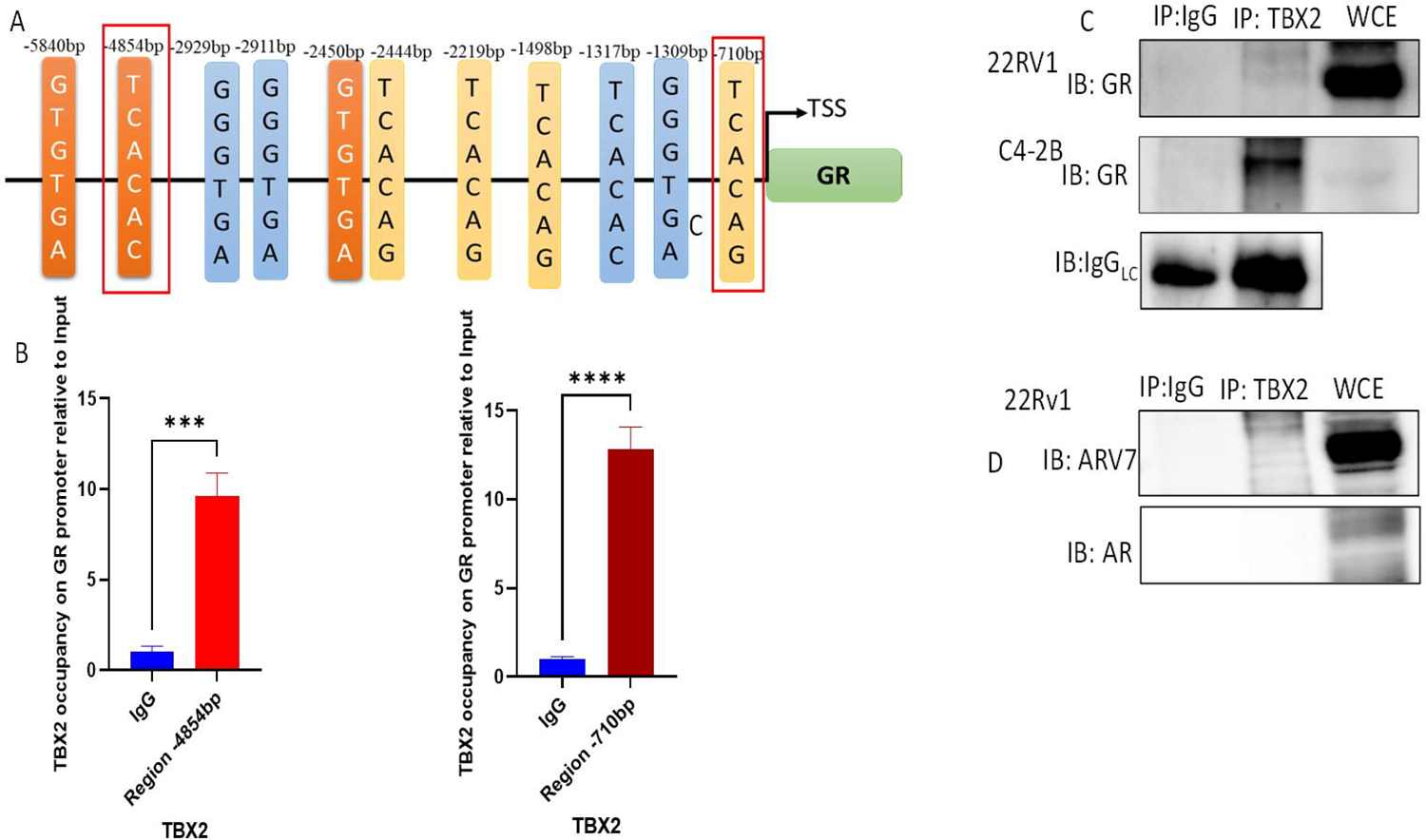
TBX2 activates GR through transcriptional activation and protein-protein interaction. **A)** *In silico* analysis of the GR promoter showing potential TBX2 binding sites; **B)** ChIP assay followed by qRT-PCR analysis showing TBX2 binding on the GR promoter with a significant enrichment of 12-fold and 8-fold respectively at −710 bp and −4854 bp upstream of the TSS; Data represent the average of triplicates ± S.D; Student’s unpaired 2-tailed *t*-tests were performed to compare the two groups **, *p*≤ 0.01; ***, *p*≤0.001; and ****,*p*≤0.0001; **C)** Western blot analysis of the CoIP in 22Rv1 and C4-2B cells showing TBX2 interaction with GR. The whole cell extract (WCE) was immunoprecipitated with an anti-TBX2 antibody with species matched IgG (control) and samples were immunoblotted for anti-GR antibody; **D)** Western blot analysis of CoIP in 22Rv1 cells showing no interaction of TBX2 with ARV7 or AR. The WCE was immunoprecipitated with an anti-TBX2 antibody with species matched IgG (control) and samples were immunoblotted for anti-ARV7 or anti-AR antibodies.

Administration of enzalutamide is currently the standard of care for advanced PCa patients since almost all patients develop resistance to the first line of ADT. The generation of AR splice variants (AR-Vs) has been attributed in large part due to the insensitivity of CRPC to enzalutamide treatment ^5^. We therefore sought to investigate if TBX2 potentially interacts physically with the ARV7 or the AR proteins. Our CoIP experiments failed to show any interaction between with ARV7 or the AR proteins (**Fig. 5D**).

### TBX2 driven switch from AR to GR signaling confers enzalutamide resistance

We next investigated if the AR to GR switch driven by TBX2 is able to confer enzalutamide resistance. Genetic modulation approaches that blocked endogenous TBX2 (PC3^TBX2DN^ and C42B^TBX2DN^) as well as the converse approach of TBX2 OE in enzalutamide sensitive LNCaP human PCa cells revealed that TBX2 expression is associated with enzalutamide resistance (**Fig. 6A**). Of note, PC3 cells have been reported to be resistant to enzalutamide treatment due to the absence of AR expression ^42, 43^, however, PC3^TBX2DN^ cells wherein AR expression is strongly upregulated - responded to enzalutamide treatment at higher doses (**Fig. 6A**). Along similar lines, LNCaP^TBX2OE^ cells displayed resistance to enzalutamide when compared with the control LNCaP^Neo^ cells (**Fig. 6A**), and as shown previously LNCaP^TBX2OE^ displayed reduced AR expression when compared with the LNCaP^Neo^ control cells (**Fig. 1D)**. Next, we investigated if knockdown of the elevated GR due to TBX2 upregulation - would result in enzalutamide sensitivity. We therefore knocked-down GR expression in LNCaP^TBX2OE^ cells which were resistant to enzalutamide treatment (**Fig. 6B, C)**. Our experiments revealed significantly decreased cell viability in enzalutamide treated LNCaP^TBX2OE^ cells in which GR was knocked-down (**Figs. 6B, C**). Taken together our results demonstrate that TBX2 driven switch from AR to GR signaling confers enzalutamide resistance since downregulation of the elevated GR in the context of increased TBX2 is sufficient to confer enzalutamide sensitivity.

**Fig. 6:**
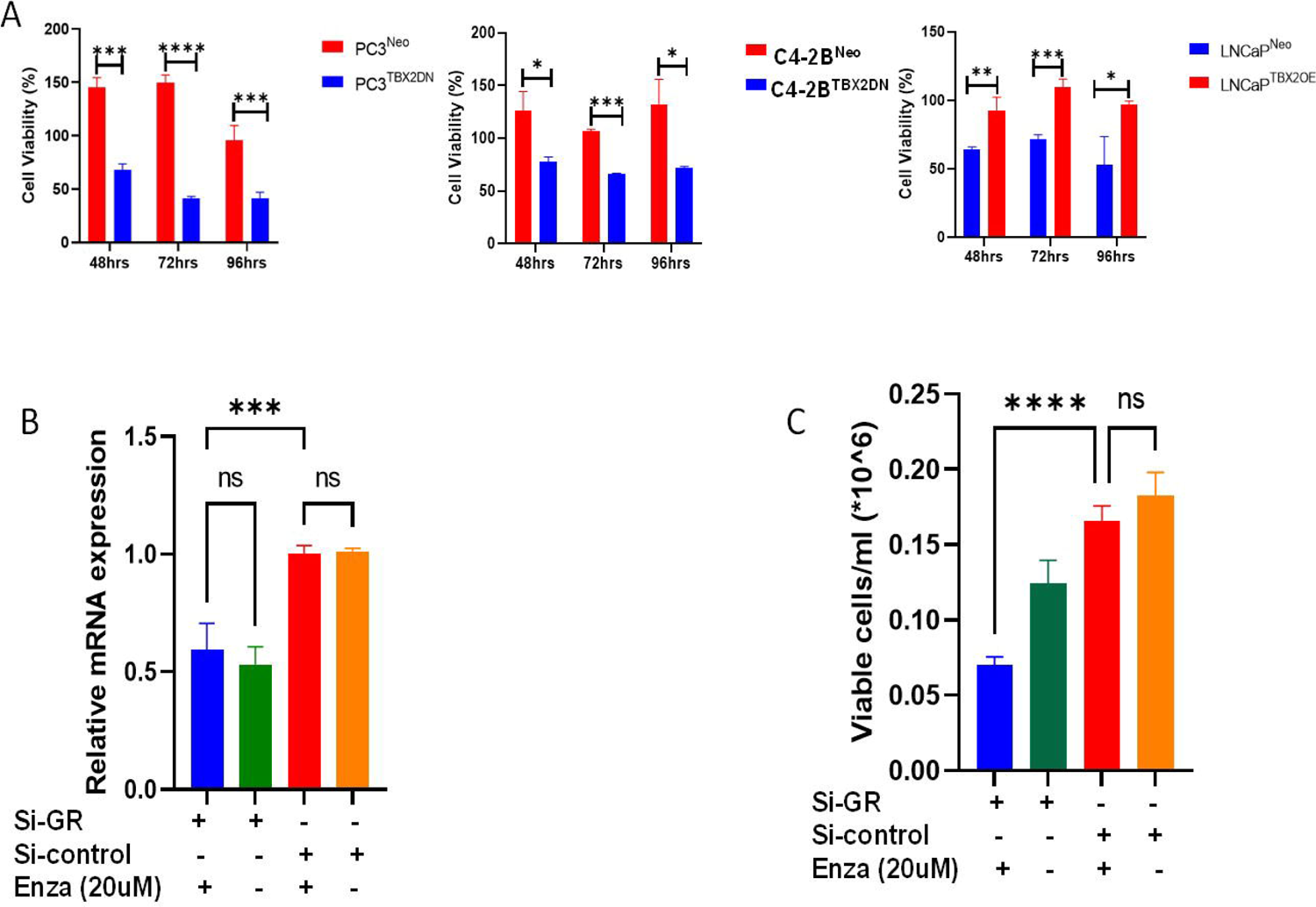
TBX2 driven switch from AR to GR signaling results in resistance to enzalutamide. **A)** Increased TBX2 expression drives enzalutamide resistance. Treatment with enzalutamide results in significant decreases in cell viability in PC3^TBX2DN^ and C4-2B^TBX2DN^ cells when compared with the respective PC3^Neo^ and C4-2B^Neo^ controls, conversely treatment with enzalutamide results in increased cell viability in LNCaP^TBX2OE^ cells when compared with LNCaP^Neo^ control cells; One way ANOVA was performed where (n=3), \*\*\**p*<0.001;****,*p*<0.0001; **B)** qRT-PCR analysis showing decreased mRNA expression of GR in LNCaP^TBX2OE^ cells that were transfected with si-GR compared with si-control with or without enzalutamide treatment; One way ANOVA was performed where (n=3), \*\*\**p*<0.001;****,*p*<0.0001; **C)** siRNA based knock-down (si-GR) of elevated GR in LNCaP^TBX2OE^ cells leads to reversal of enzalutamide resistance to enzalutamide sensitivity. Upon treatment with enzalutamide, significant decrease in cell viability is observed in LNCaP^TBX2OE^ cells that were transfected with si-GR (blue bar) when compared with si-control (red bar); One way ANOVA was performed where (n=3), \*\*\**p*<0.001;****,*p*<0.0001.

### SP-2509, an allosteric LSD1 inhibitor, impedes TBX2-GR physical interaction

Having identified GR as a novel interactor of TBX2 in CRPC, we next sought to investigate the possibility of pharmacologically targeting the TBX2-GR interaction. Previous studies have identified an interaction between TBX2 and LSD1 as a part of the COREST complex ^17^. These reports also identified that the interaction between TBX2 and LSD1 could be pharmacologically targeted through the LSD1 inhibitor SP2509 ^17^, and that disruption of the TBX2-LSD1 interaction resulted in the inhibition of TBX2 downstream targets ^17^. Further, and of relevance to our study, another report showed that LSD1 is upregulated in CRPC ^44, 45^. We therefore sought to determine if SP2509 had the ability to disrupt TBX2-GR interaction. Following the determination of the optimum dose of SP2509 dose, we performed a CoIP in SP2509 treated 22Rv1 and C4-2B human CRPC cells wherein we observed that SP2509 disrupted the interaction between TBX2 and GR in addition to disrupting LSD1-TBX2 interaction as reported previously ^17^ (**Figs. 7A, B, C, D).** These results provide vital proof-of-concept insights about the potential of pharmacologically inhibiting TBX2-GR interaction, and thereby target the TBX2-driven switch from AR to GR signaling in CRPC.

**Fig. 7:**
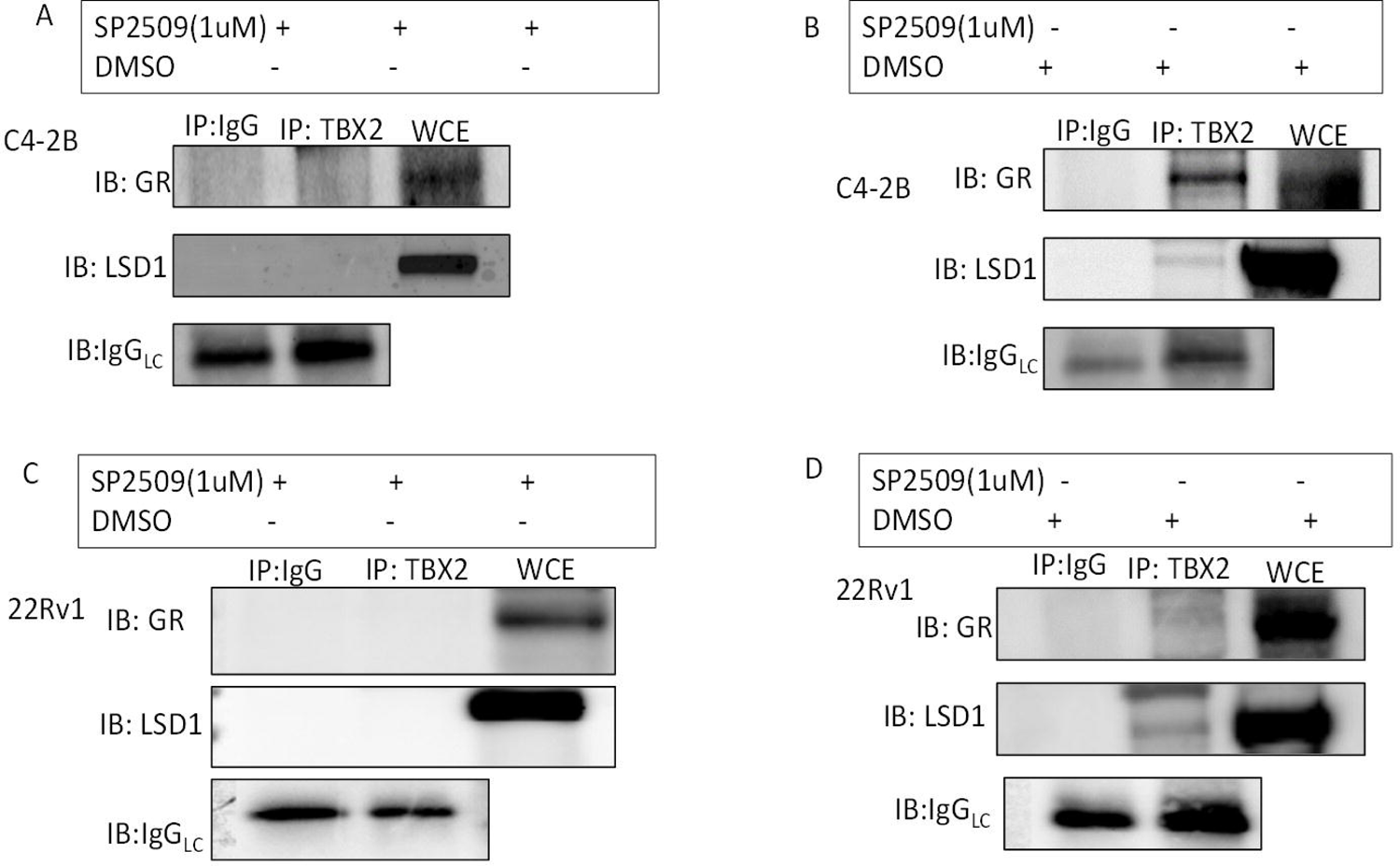
SP2509, an allosteric LSD1 inhibitor disrupts TBX2-GR protein-protein interaction. **Fig. 7**: SP2509, an allosteric LSD1 inhibitor disrupts TBX2-GR protein-protein interaction. A) Western blot analysis of the CoIP in SP2509 treated C4-2B cells shows loss of TBX2-GR and TBX2-LSD1 interactions. Samples were immunoprecipitated with an anti-TBX2 antibody using species matched control IgG and immunoblotted for anti-LSD1 or anti-GR antibodies; B) Western blot analysis of the CoIP in C4-2B cells treated with DMSO vehicle control showing TBX2-GR and TBX2-LSD1 interactions. The whole cell extract (WCE) was immunoprecipitated with an anti-TBX2 antibody with species matched control IgG and samples were immunoblotted for anti-LSD1 or anti-GR antibodies; C) Western blot of CoIP in SP2509 treated 22Rv1 cells shows loss of TBX2-GR and TBX2-LSD1 interactions. The WCE from 22Rv1 cells was immunoprecipitated with an anti-TBX2 antibody using species matched IgG as control and immunoblotted for anti-LSD1 or anti-GR antibodies; D) Western blot of CoIP in 22Rv1 cells treated with DMSO vehicle control showing TBX2-GR and TBX2-LSD1 interactions. WCE from 22Rv1 was immunoprecipitated with an anti-TBX2 antibody using species matched IgG as control and samples were immunoblotted for anti-LSD1 and anti-GR antibodies.

## Discussion

Recent evidence has demonstrated that acquired-resistance to ADT is often due to upregulated expression of the GR ^7–11, 13^. The GR, a steroid nuclear receptor akin to the AR, shares a significant homology to AR including an overlapping target gene set ^9, 10, 13, 15, 16^. This has led to the hypothesis that the 2^nd^ generation ARSIs, such as enzalutamide, put selective pressures on PCa cells that drive the clinically unintended consequence of AR to the GR signaling bypass ^7–11^. Previous reports have shown that GR is negatively regulated by the AR ^9, 46, 47^. However, fundamental unanswered questions remain, including the identification of the specific molecular mechanisms that drive the AR to GR signaling bypass, and why the steroid receptor signaling bypass results in a generalized failure of current therapeutic modalities. Also, can indirect modes of GR signaling blockade ^11, 48^ be identified that offer therapeutic efficacy while avoiding the serious collateral toxicities of systemic GR signaling inhibition ^12, 22^

In an effort to identify drivers of lethal disease progression in PCa, our previous study that utilized various xenograft mouse models, was the first to report that TBX2, a developmental T-box transcription factor (TF) master regulator is over-expressed in CRPC ^18^ and drives bone metastatic progression in PCa ^18^. In agreement with our findings, a recent report confirmed that TBX2 and GR were two of the four TFs that drive enzalutamide resistance in advanced PCa ^21^.

Our first-in-field observations show that TBX2 is the molecular switch that represses AR while activating GR to drive the emergence of CRPC (**Fig. 8A)**. Specifically, we detected a negative correlation between TBX2 and AR expression, and a positive correlation between TBX2 and GR expression, in publicly available PCa clinical datasets. Then, using multiple complementary approaches to genetically modulate TBX2 expression in human PCa cells *in vitro*, we determined that TBX2 down-regulates the AR, and up-regulates the GR. Mechanistically, our studies revealed that TBX2 repressed AR and GATA2 by direct binding to the promoters; and conversely TBX2 upregulated the GR through a dual-mechanism of: a) direct binding of TBX2 to the GR promoter, and b) TBX2-GR protein-protein interaction. Further, using genetic modulation approaches for TBX2, our studies demonstrated that TBX2 driven switch from AR to GR signaling confers enzalutamide resistance in CRPC.

**Fig. 8:**
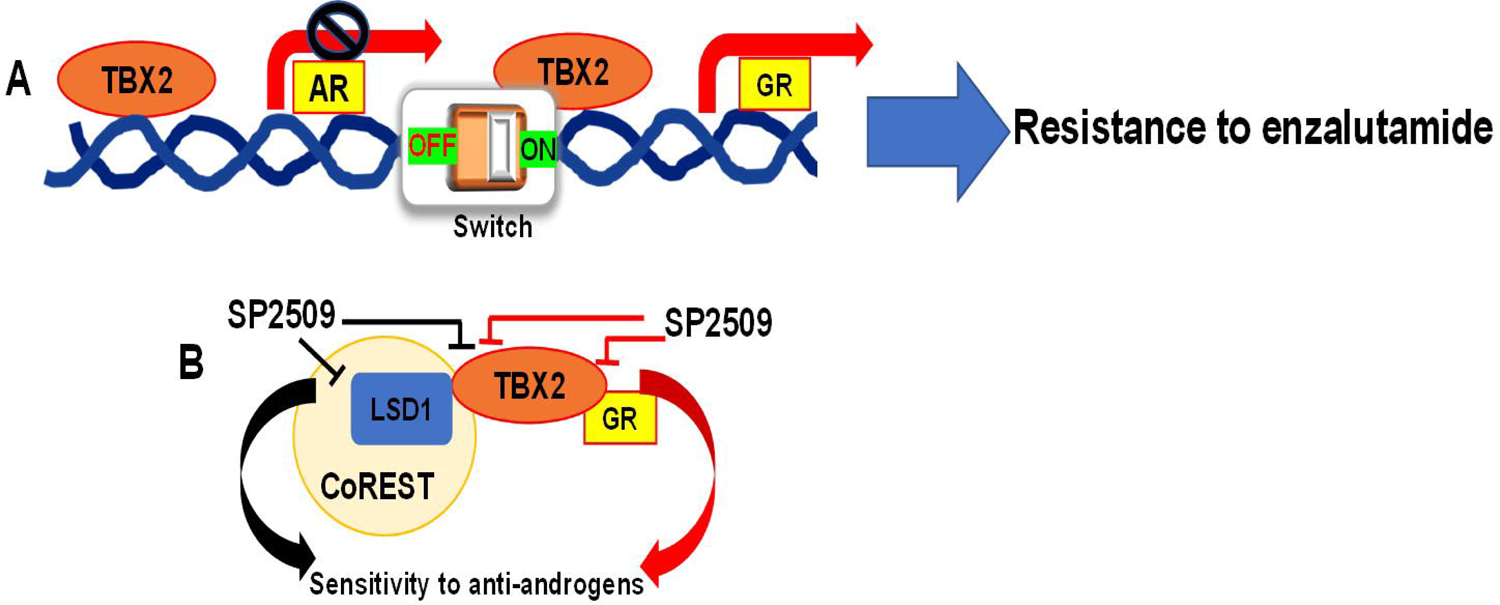
TBX2 mediated switch from Androgen Receptor to Glucocorticoid Receptor signaling drives therapeutic resistance in Prostate Cancer. A) TBX2 acts as a switch to turn OFF AR on one hand and turn ON GR on the other to orchestrate resistance to enzalutamide in CRPC; B) SP2509, an allosteric inhibition of LSD1 could be utilized to disrupt both TBX2-LSD1 (previously reported and represented as a black block-arrow) and TBX2-GR protein-protein (based on our findings, and represented as a red block-arrow) interactions, and thereby reinstating sensitivity to anti-androgens.

TFs, including TBX2, have traditionally been considered “undruggable” targets ^49^. However, recent reports ^17^ have highlighted that TBX2 target specificity is dependent to a larger extent on its protein-protein interactions providing crucial insights into therapeutic strategies for targeting TBX2 via disruption of its protein partners. These studies showed that TBX2 forms a complex with lysine-specific demethylase 1 (LSD1) ^17^, a protein who’s demethylase independent function is associated with CRPC ^45^. In addition, these reports demonstrated that allosteric inhibition of LSD1 via SP2509 in addition to disrupting TBX2-LSD1 interaction de-represses TBX2 targets, pointing to the fact that the target specificity of TBX2 is dictated by its protein partners ^17^. Along the same lines, these studies showed that silencing of LSD1 and its protein partners including TBX2 emulates the effect of TBX2 knock-down ^17^ resulting in de-repression of TBX2 targets. Importantly, LSD1 inhibitors including SP2509 are currently in clinical trials for treatment of various cancers thereby making SP2509 a potential therapy for PCa ^50^.

The function of SP2509 as an allosteric inhibitor of LSD1 seems to be highly relevant to the CRPC phenotype. This is borne by the fact that LSD1’s role in driving CRPC is independent of its function as an AR co-activator^51^ which is dependent on its canonical histone demethylase activity ^45^. In accordance, these reports demonstrated that SP2509 can disrupt the interaction of between LSD1 and its protein partners resulting in enzalutamide sensitivity while not altering its histone demethylase function ^17^. Intriguingly, TBX2 function in driving the CRPC phenotype parallels the ability of LSD1 and its binding partners in mediating CRPC ^44, 45^. The enhanced stemness phenotype in relation to LSD1’s function in CRPC ^44, 45^ is in consonance with our previous report on TBX2 wherein we found that TBX2 upregulates SOX2 and MYCN in CRPC both via autocrine and paracrine modes ^19^.

Our finding that GR physically interacts with TBX2, juxtaposed with our observation that SP2509 can disrupt both TBX2-GR and TBX2-LSD1 interactions (**Fig. 8B)**, raises the possibility that GR is a part of the COREST complex. However, further details including the other protein interactors of the GR within the complex remain to be elucidated. Of note, our findings that TBX2 acts as the molecular switch that drives the AR to GR signaling bypass in conjunction with our model that TBX2 and GR could be part of the COREST complex provides novel implications for the use of SP2509 in the treatment of enzalutamide resistant CRPC. Allosteric inhibition of LSD1 has previously been reported to confer enzalutamide sensitivity ^45^. However, our findings that TBX2 and GR as a part of a protein complex point to a model wherein allosteric inhibition of LSD1 could additionally result in the disruption of TBX2-GR interaction. Taken together our findings on the TBX2/GR signaling shed light on a hitherto unknown function of SP2509 in conferring sensitivity to enzalutamide i.e. via reversing the AR to GR signaling bypass (**Fig. 8B)**. GR ablation approaches have deleterious consequences and have been unsuccessful in the treatment of PCa ^12, 22^. Therefore, our proposed model (**Figs. 8A, B)** wherein GR could be inhibited in an indirect manner in CRPC via targeting the upstream AR to GR molecular switch that is driven by TBX2 may provide a potential therapeutic strategy for preventing acquired resistance in the lethal progression of the disease.

## Authors’ Contributions

**Conception and design:** S.D., G.K.P., M.T., S.N.

**Development of methodology:** S.D., G.K.P., M.T., S.N.

**Acquisition of data (provided animals, acquired and managed patients, provided facilities, etc.):** S.D., G.K.P, H.K, D.L., M.T., S.N.

**Analysis and interpretation of data (e.g., statistical analysis, biostatistics, computational analysis):** S.D., G.K.P, M.T., S.N.

**Writing, review, and/or revision of the manuscript:** S.D., M.T., S.N.

**Study supervision:** M.T., S.N.

**Administrative, technical, or material support (i.e., reporting or organizing data, constructing databases):** S. D., G.K.P., H.K., M.T., S.N.

## Acknowledgments

The authors would like to thank Robert J. Matusik (Vanderbilt University Medical Center) for critical reading of the manuscript. The authors acknowledge Colin Goding (Ludwig Cancer Research, University of Oxford, United Kingdom) and Maarten van Lohuizen (NKI-AVL, Amsterdam, the Netherlands) for their generous gift of viral vectors namely, p-BABE-puro-TBX2DN and LZRS-TBX2-iresGFP constructs, respectively. The authors would like to acknowledge Sambantham Shanmugam for help in the luciferase reporter assay.

